# Automated multiscale high-content imaging connects plaque population heterogeneity to localized synapse loss

**DOI:** 10.64898/2026.10.26.761378

**Authors:** Celia A. Polino, Russell Port, Mackenzie Hofmann, Sean Smith, Xiaohai Wang, Rebecca Klein, Michael Marino

**Affiliations:** Neuroscience Discovery, Merck & Co., Inc., West Point, Pennsylvania, USA

**Keywords:** High-content imaging, pathology, synapse, Alzheimer’s Disease, amyloid beta, automation

## Abstract

Synapse loss is a major correlate of cognitive decline in neurodegenerative disease, yet its relationship to localized pathology remains difficult to study at scale. Conventional confocal imaging approaches are limited by low throughput, selection bias, and restricted sampling of pathological features. Here, we describe an automated high-content imaging and analysis workflow that enables large-scale quantification of amyloid beta plaque-associated synapse loss within mouse brain tissue. Using Preciscan-based multiscale acquisition on the Opera Phenix platform, this workflow combines tissue-wide pathology mapping with targeted high-magnification imaging and subcellular analysis. We apply this framework to acquire and analyze hundreds of plaque images in the CRND8 mouse model of Aβ proteinopathy, demonstrating plaque-associated synapse loss within developing plaque populations over disease progression. We find that small diffuse plaques accumulate rapidly at late stages in CRND8, but that the primary synapse loss comes from large cored plaques. More broadly, this scalable framework enables systematic investigation of subcellular phenomena across pathological microenvironments in disease models.

## Introduction

Loss of synaptic integrity is strongly associated with cognitive decline and disease progression in neurodegenerative diseases, reflecting neuronal dysfunction and the impact of pathological processes^1–3^. Importantly, synaptic alterations often occur early in disease and are spatially heterogeneous, with synapses showing selective vulnerability in relation to local pathological features^4,5^. For example, excitatory synapses show pronounced, distance-dependent loss in the immediate vicinity of amyloid beta (Aβ) plaques, with relatively preserved density at greater distances^6–9^. In addition, synapse loss across tau-rich regions follows patterns of local tau accumulation and network connectivity, and is more associated with neuritic tau pathology than with neurofibrillary tangles^4,10,11^. Together, these findings indicate that synaptic vulnerability is in large part governed by specific, spatially-restricted microenvironments, rather than by a brain-wide disease state.

Yet across studies employing human tissue and animal models, synaptic density is typically imaged and quantified within a limited number of manually selected fields or brain regions ^6,8,9,11–23^. These approaches introduce several key limitations. Critically, manual workflows require substantial hands-on effort for field selection, image acquisition, and analysis, limiting scalability and severely limiting throughput and sample size. In addition, manual field selection may fail to capture specific pathologies of interest, leading to potential sampling bias and loss of spatial context. As a result of these constraints, many researchers reduce experimental burden by turning to surrogate readouts such as total synaptic protein levels within brain homogenates or synaptosome-enriched fractions^13,24–27^. However, analyses of homogenized tissue dilute localized synaptic signals and eliminate spatial information, preventing analysis of how synapses relate to pathological features, what markers correlate with localized synapse loss, and how best to assess these processes in human patients. Together, these approaches emphasize global synaptic changes but lack the capabilities required to quantify synapse alterations within the microenvironments where pathology and therapeutic effects are often localized.

This fundamental trade-off between high spatial resolution and tissue-wide analysis is not restricted to studies of pathology-associated synapse loss. In the field of neurodegeneration, subcellular mechanisms such as the functions of mitochondria ^28,29^, lysosomes ^30,31^, and lipid droplets ^32–34^ are emerging as key influences on neuropathology and are increasingly a focus of therapeutic development. Elucidating these mechanisms requires bridging subcellular resolution with macro-scale anatomical context to fully understand how these processes work in disease. This highlights a major technical gap and an opportunity for automated, pathology-associated imaging approaches.

Rather than relying on manual, user-directed selection of representative fields, emerging imaging technologies can enable unbiased identification of pathological features or cell populations of interest, cohort-wide throughput, and automated downstream analysis, all while preserving subcellular resolution. These advances allow researchers to move beyond sparsely sampled, potentially biased datasets toward systematic, large-scale characterizations of complex phenomena. By streamlining and standardizing data acquisition, such workflows significantly reduce acquisition and analysis time while improving reproducibility across experiments and cohorts.

Here we describe a hierarchical pipeline for the Opera Phenix with automated Aβ plaque detection and selection of 20× fields for re-imaging at 63× magnification. Using this workflow, we can collect hundreds of highly comparable plaque images across dozens of mouse brains per slide, with sufficient resolution to quantify distance-based synapse loss and its relationship to plaque morphology and animal age. We apply this framework across a time course in the CRND8 amyloid beta plaque mouse model to illustrate how plaque-associated features evolve alongside progressive synapse loss.

We envision that similar workflows can be adapted to other imaging platforms to greatly enhance the spatial resolution and throughput of subcellular analyses, enabling systematic characterization of pathological microenvironments and cellular interactions that are otherwise difficult to capture at scale.

## Results

### Automated multiscale high-content imaging enables plaque-targeted subcellular analysis across multi-brain block sections

To enable systematic analysis of pathology-associated subcellular features, we established a multiscale high-content imaging workflow using the Opera Phenix platform with Preciscan hierarchical acquisition (Fig. 1). This method combines the capabilities of the Opera Phenix spinning disc high-content imager, multi-brain tissue blocks from Neuroscience Associates (NSA) labs, and a custom 3D-printed slide holder created in-house (Fig. 1A). The Harmony “Preciscan” feature of the Opera Phenix is designed to “Prescan” low-magnification fields, analyze them to assign a new well layout based on regions of interest, and “Rescan” at a higher magnification. We have applied this method to characterize plaque-associated synapse loss, utilizing multiplex immunofluorescence consisting of 6E10 to detect total Aβ, Methoxy-X04 (Methoxy) to identify fibrillar plaque cores, LAMP2 to visualize plaque-associated dystrophic neurites (DNs), and HOMER1 to quantify postsynaptic puncta. Cortex regions were chosen for the “Prescan” as the cortex reliably accumulates Aβ plaques in the CRND8 model and has a relatively homogenous neuropil for synaptic quantification. Prescans consisted of 20× tile-scanned z-stacks to generate a low-magnification image of pathology across the entire multi-brain block section (Fig. 1B). Preciscan analysis of the 20× scan was then used to identify and filter plaques based on several features, which informed the subsequent automated “Rescan” of selected fields with a 63× water objective. This hierarchical acquisition strategy preserves tissue-scale information while providing the resolution required to visualize individual synaptic puncta in relation to plaque pathology.

**Figure 1.**
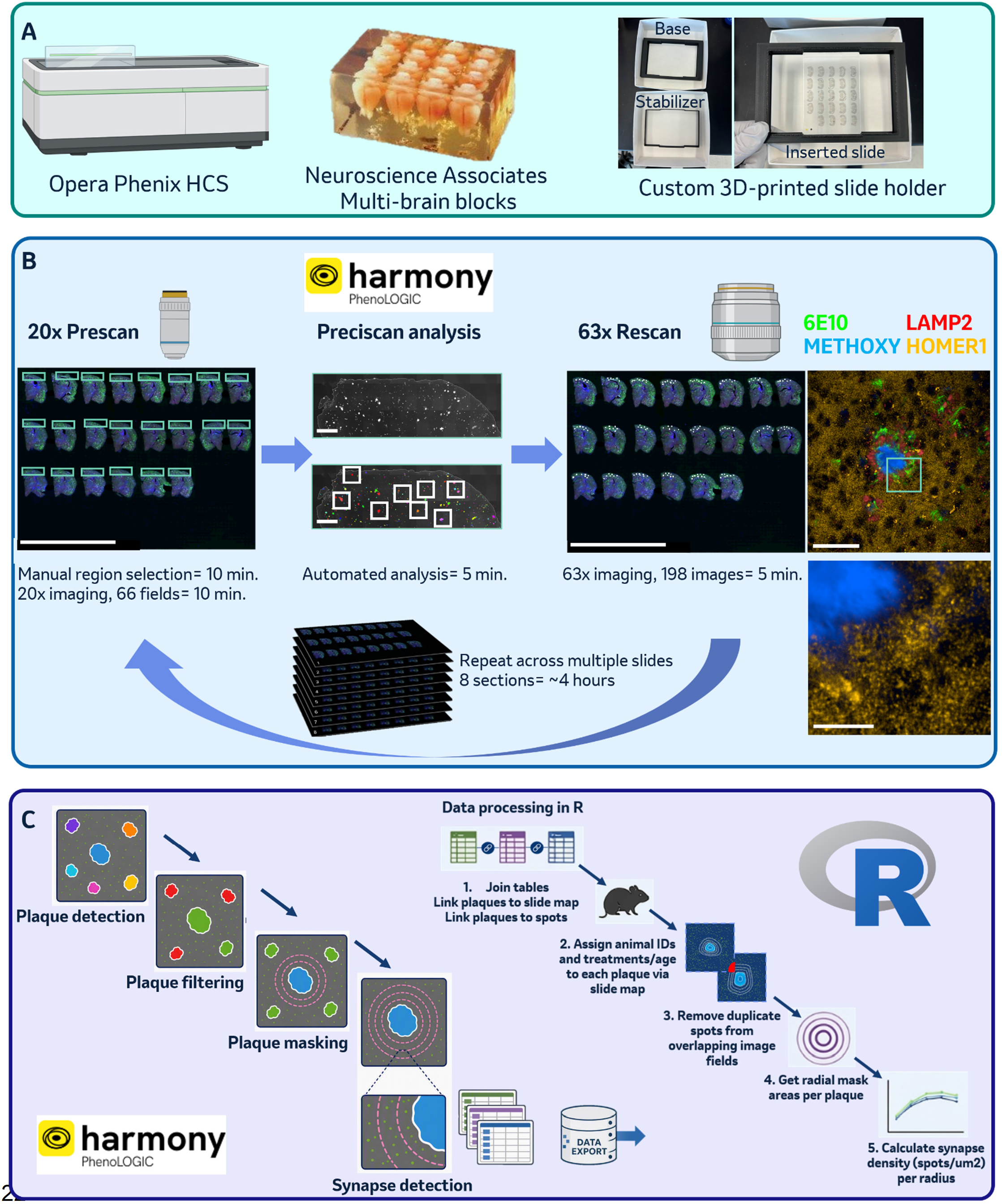
High-Content Imaging with the Opera Phenix Preciscan enables imaging subcellular features across multi-brain block sections. **(A)** Imaging tools, including the Opera Phenix high-content screening system, multi-brain tissue blocks from NSA labs, and a custom 3D-printed slide holder consisting of a base and stabilizer. **(B)** Hierarchical acquisition workflow. Cortical regions (teal boxes) from multiple brains were manually selected and tile-scanned at 20× magnification (Scale bar, 20mm). Preciscan analysis was then used to identify plaques within each cortex (white boxes) for automated re-imaging with a 63× objective (Scale bars, 500µm). In the representative workflow, manual selection of 22 cortical regions required approximately 10 min, 20× imaging of three fields per brain required approximately 10 min, and automated Preciscan analysis required approximately 5 min. Subsequent 63× imaging of up to three plaque fields (white boxes) per 20× field required approximately 5 min and enabled acquisition of up to 198 high-magnification plaque images across 22 brains per block section. Repetition across multiple slides enabled imaging of eight block sections in approximately 4 h. Representative 63× plaque image and high-magnification inset shows plaque-associated synaptic puncta (63× plaque image scale bar, 50µm, inset image scale bar, 12.5 µm). **(C)** Image-processing and data-analysis workflow. Plaques were detected and filtered in Harmony, plaque-associated concentric masks were generated and used for detection of HOMER1 synaptic puncta. Slide-level, plaque-level, and puncta-level outputs were linked in R using slide-map X/Y coordinates, duplicate synapse counts from overlapping image fields were removed, and synapse density was calculated within the plaque and surrounding concentric ring masks.

This workflow substantially decreases the time to collect hundreds of high-magnification plaque images. In the representative acquisition shown, manual selection of 22 cortical regions and 20× Prescan imaging each required approximately 10 minutes, while automated Preciscan analysis required approximately 5 minutes (Fig. 1B). 63× imaging of up to 3 plaque fields per each 20× field enabled a balance of optimal plaque sampling with minimal image overlap, and required approximately 5 minutes to generate as many as 198 high-resolution plaque images across 22 mouse brains. Repetition of this process across multiple slides enabled acquisition from eight block sections in approximately four hours, demonstrating the feasibility of extending subcellular imaging across large tissue cohorts rather than relying on a limited number of manually selected fields.

Following image acquisition, a Harmony and R analysis pipeline enabled filtering of plaques, detection of synapses, and wrangling of data from thousands of synapses to give per-plaque, per-animal, and per-group quantification (Fig. 1C). In Harmony, plaques were detected, filtered, and expanded into spatial masks for quantification of plaque-associated synaptic puncta. The Harmony-exported slide, plaque, and puncta-level tables were joined in R using slide-map X/Y position, allowing each plaque and surrounding synapses to be associated with the corresponding animal and experimental group. Overlapping fields were reconciled to remove duplicate synapse measurements, and concentric radial masks were used to calculate synapse density as a function of distance from each plaque. Together, this workflow generates plaque-associated synapse measurements that retain the connection between local plaque characteristics and tissue-wide pathology.

### Automated plaque detection, filtering, and field selection with Preciscan links 20× Prescan imaging to targeted 63× Rescan acquisition

To achieve automated plaque-targeted imaging, 20× Prescan acquisitions were analyzed to identify candidate plaques and determine their optimal X/Y/Z coordinates for subsequent high-magnification imaging (Fig. 2A). Prescan datasets were acquired as tiled 20× z-stacks to enable focal plane selection. Prescan images were assembled into global images and corresponding plane maps that recorded optimal focal planes across the multi-brain block section. These plane maps provided a representation of section topography and were used to guide downstream image acquisition.

**Figure 2.**
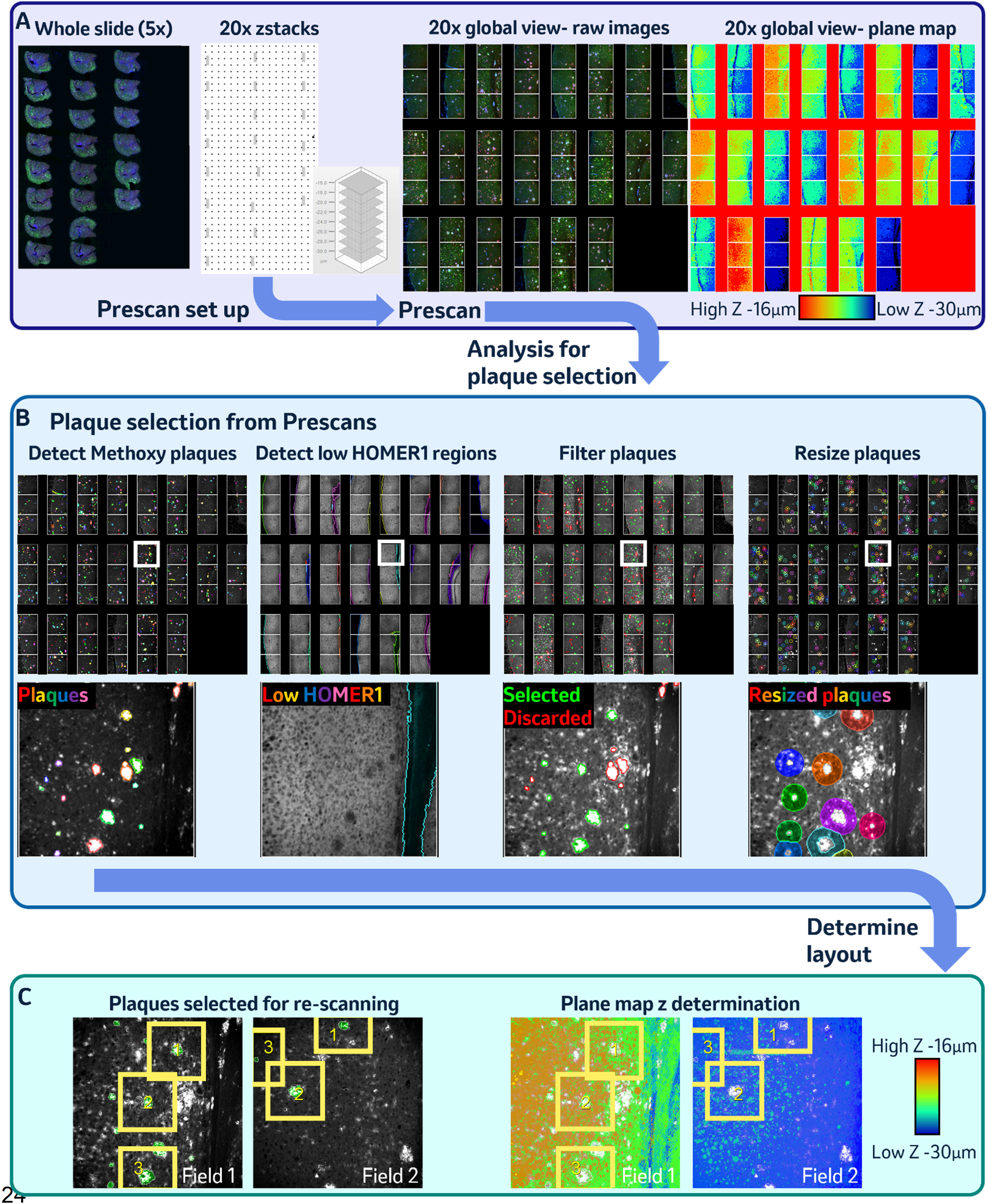
Opera Phenix “Preciscan” analysis links 20× Prescan imaging to analysis and filtering for 63× Rescan imaging. **(A)** Prescan setup and acquisition. Cortex images from multi-brain slides were acquired as tiled 20× z-stacks. The individual images were assembled into a global raw-image view and a corresponding plane map representing the selected focal position across each tissue region. Plane-map colors indicate relative z position, ranging from approximately -16 µm at the highest positions to -30 µm at the lowest positions in the representative acquisition. **(B)** Prescan analysis sequence for selection of plaque-centered images. Methoxy+ plaque objects were detected. Regions with low HOMER1 signal were identified for exclusion (usually the corpus callosum or outside of the tissue). Candidate plaques were filtered on size and lack of close neighbors to identify unsuitable targets. Selected plaque objects were resized to measure HOMER1 intensity within an extended ring region. **(C)** Generation of the Rescan layout. Selected plaques were assigned to 63× imaging fields, with numbered boxes indicating individual plaque targets grouped within a 20× field. The plane z-map shows variation in the focal plane between 20× fields and across an individual field. Thus, z-planes for the Rescan were assigned per field based on the HOMER1 intensity in the resized plaque ring region.

For the example dataset we describe below, we chose to focus imaging on Methoxy-X04 (Methoxy+) plaques. Our preliminary imaging tests showed that Methoxy-negative (Methoxy-) diffuse plaques were frequent enough that they would still be included alongside Methoxy+ plaques in Methoxy-centered images (Fig. 6C). Methoxy+ plaques were detected throughout the Prescan images and subjected to a series of automated filtering steps (Fig. 2B). Regions lacking sufficient HOMER1 signal (usually the corpus callosum or outside of the tissue) were identified and excluded from analysis. Plaques were then filtered according to size (minimum 200 µm^2^, maximum 2500 µm^2^) and presence of close neighbors (to reduce selection of plaques with overlapping masks). Selected plaques were subsequently resized to check the surrounding tissue for adequate HOMER1 signal. The Z coordinate of each selected resized plaque was determined from the Prescan plane map, allowing Preciscan to automatically determine the appropriate focal plane for each single-plane 63× image field (Fig. 2C). This process connected low-magnification pathology detection with automated high-magnification imaging, enabling efficient acquisition of plaque-centered fields and avoiding manual selection of individual plaques or imaging locations.

As shown in Supplemental Figure 1, a similar workflow was used to detect all Methoxy, 6E10, and dystrophic neurite objects in order to analyze cortex-wide pathological populations; however, filters intended to select isolated plaques suitable for synapse quantification, including maximum size, nearest-neighbor, and low-HOMER1 exclusion criteria, were not applied. A minimum size threshold was retained to exclude small debris.

### Post-Rescan filtering identifies plaques suitable for quantitative synapse analysis

While Preciscan acquisition successfully generated large numbers of plaque-centered images, additional post-imaging filtering was required to ensure selection of plaques suitable for quantitative synapse analysis. Some Rescan images were not ideal for synapse quantification for varying reasons, such as poor HOMER1 signal due to inaccurate z-plane selection, close neighboring plaques with overlapping mask regions not apparent at low-magnification, etc. Thus, several filters were applied to Rescan images with the full flexibility of the Harmony analysis software in order to control for plaques with ideal features.

In the example shown in Figure 3, following acquisition of 176 high-resolution plaque-centered images across one block section, intensity thresholding of the 6E10 channel identified 739 candidate 6E10-positive objects across the dataset. To distinguish true plaque structures from non-plaque objects, a PhenoLOGIC machine-learning classifier was trained across multiple Rescans and applied to the dataset. This step enabled exclusion of objects with plaque-level 6E10 intensity that lacked plaque characteristics, including occasional blood vessel-associated signal. Objects classified as plaques were also filtered using minimum-size criteria to remove small detections unlikely to represent biologically relevant Aβ deposits.

**Figure 3.**
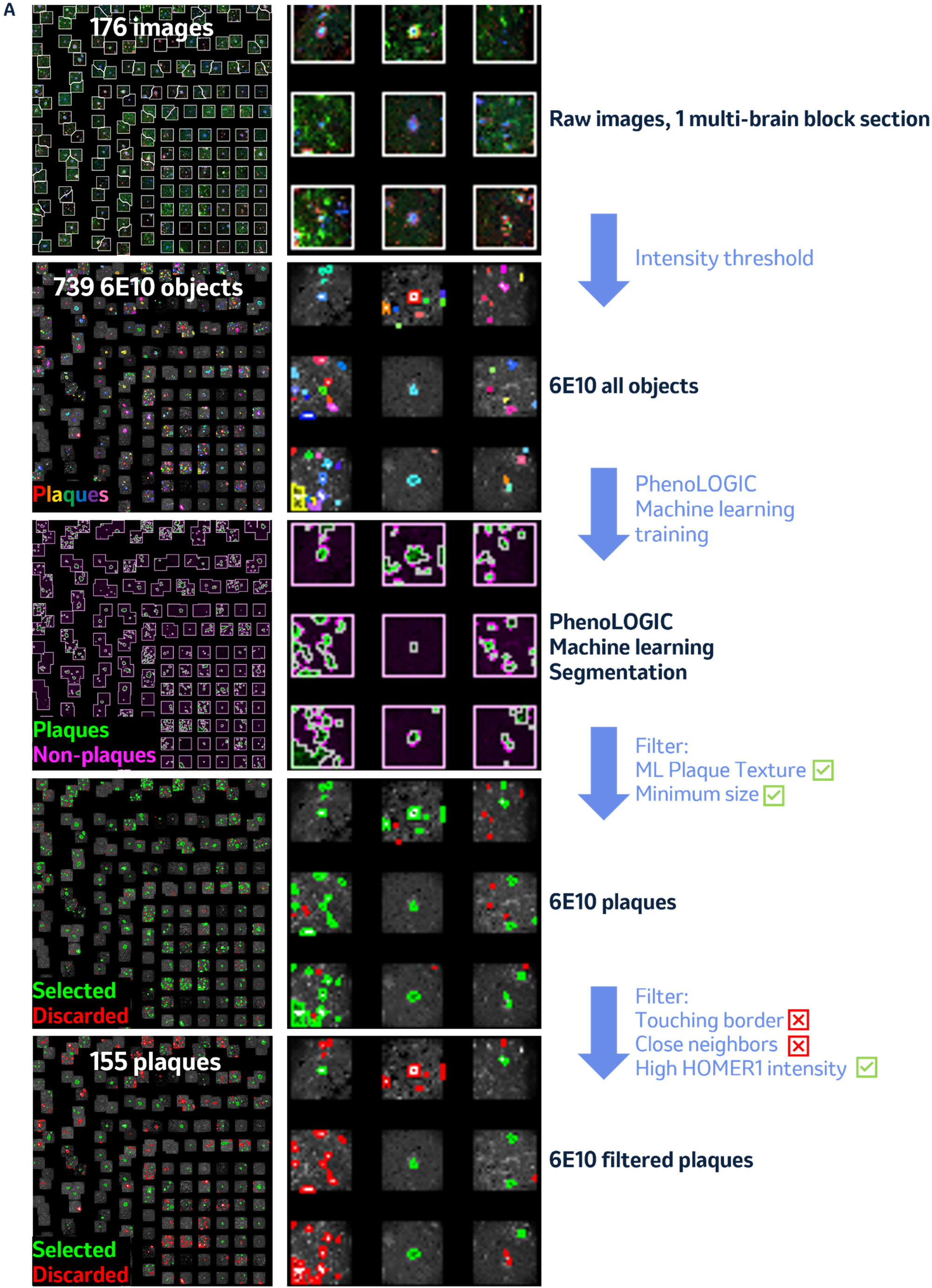
Post-imaging filtering allows selection of individual plaques with ideal features for synapse quantification. **(A**) Representative post-acquisition analysis workflow applied to 63× plaque-centered images. Whole-dataset views are shown at left, and representative inset images with segmentation overlays are shown at right. The example dataset from all brains on one block section contained 176 images. Following intensity thresholding, 739 6E10+ objects were detected across all images. PhenoLOGIC machine-learning segmentation was trained to distinguish plaques from non-plaque objects. This enabled exclusion of blood vessels which may be faintly 6E10+. This machine-learning plaque texture along with a minimal size filter was used to classify objects as 6E10 plaques. Additional quality-control filters excluded plaques touching an image border or located near neighboring plaques. Plaques were also filtered to have a high surrounding HOMER1 intensity in order to ensure optimal puncta quantification for synapse density. For this example block section, the final filtered dataset contained 155 plaques of mixed size and morphology.

Additional quality-control filters were then applied to optimize subsequent synapse quantification. Plaques touching image borders were excluded because the complete plaque perimeter and surrounding tissue could not be analyzed. Plaques located near neighboring plaques were removed to prevent overlap between plaque-associated mask regions. Finally, plaques were required to exhibit sufficient surrounding HOMER1 signal to ensure reliable synaptic puncta detection within plaque-associated masks.

Application of these sequential filters reduced the dataset from 739 candidate 6E10-positive objects to 155 plaques suitable for synapse analysis. This strict filtering strategy generated a dataset composed of isolated plaques with well-defined borders and quantifiable surrounding synaptic signal.

### Age-dependent progression of plaque pathology in CRND8 mice is driven predominantly by the accumulation of small diffuse plaques

To apply this multiscale imaging workflow to a relevant disease model, we performed a time-course study of the CRND8 model by collecting brains from mice at 6, 8, 10 and 12 months of age. As described above, brains were stained with a combination of Methoxy-X04 for fibrillar plaque cores, 6E10 for total Aβ, LAMP2 for dystrophic neurites, and HOMER1 for post-synaptic puncta. This staining set combined with 20×-63× Preciscan imaging allowed us to quantify tissue-wide plaque populations in 20× images, re-image plaques and synapses at 63×, and combine these observations in order to understand how synapse loss measured at individual plaques may affect the overall tissue.

We first quantified 6E10+ plaques, Methoxy+ plaques, and LAMP2+ dystrophic neurite populations imaged across the cortex at 20× for 8 block sections (Fig. 4A). We plotted size distribution histograms for all objects both by object frequency and percent tissue area (Fig. 4B) and analyzed the data with a mixed-effect model followed by Tukey’s multiple comparisons (Supp. Fig. 2). This allowed us to test whether objects at each size bin differed based on animal age.

**Figure 4.**
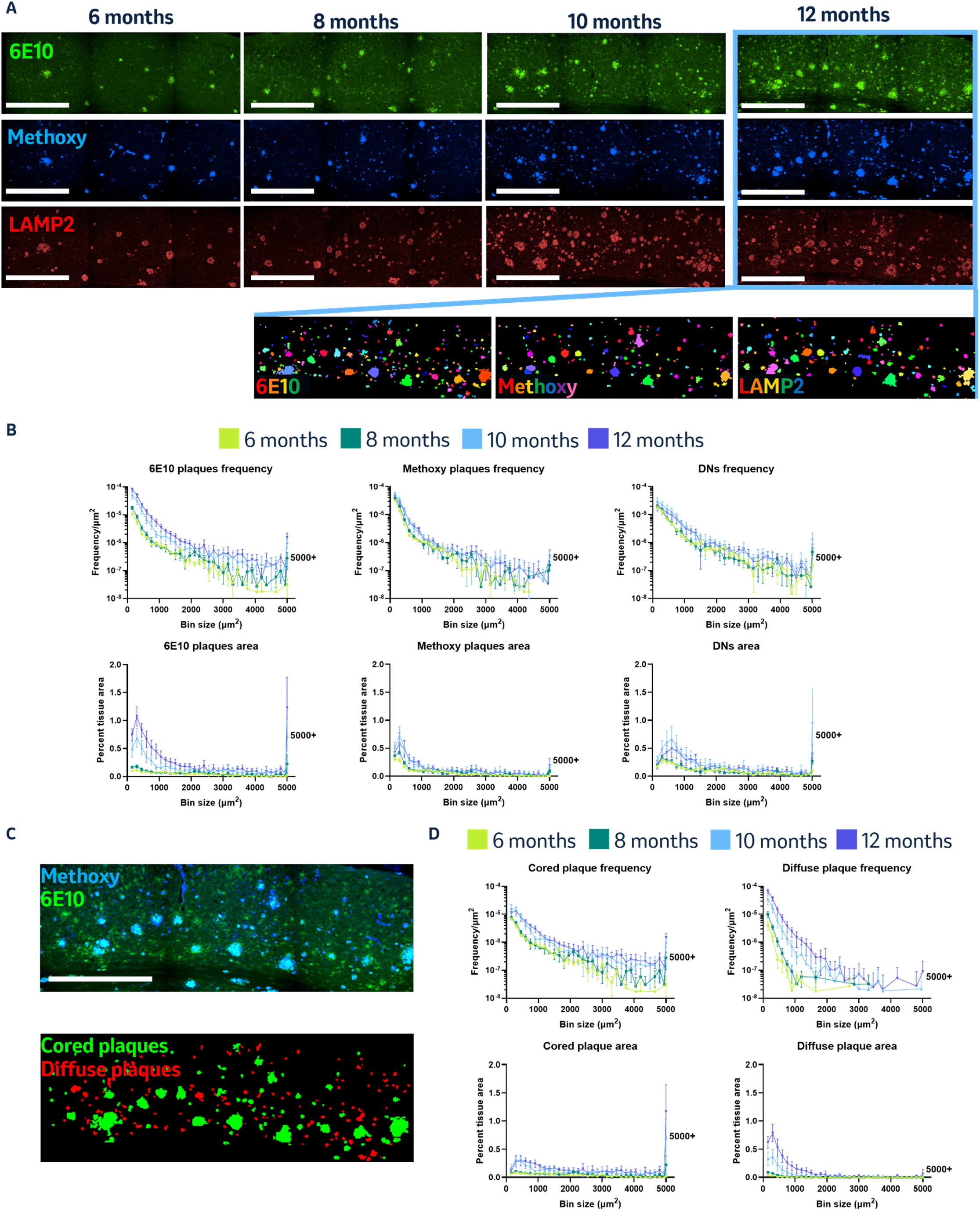
Cored plaques accumulate early and diffuse plaques continue to develop at late stages in the CRND8 model. **(A)** Representative 20× cortical images of 6E10, Methoxy, and LAMP2 dystrophic neurites (DNs) from CRND8 mice at 6, 8, 10, and 12 months of age (Scale bars, 500µm). The inset images indicate object segmentation for each stain from a 12-month cortex. **(B)** Size distributions of 6E10+ plaques, Methoxy+ plaques, and DNs at each age. The upper plots show object frequency across size bins, and the lower plots show the percentage of analyzed tissue area occupied by objects within each size bin. **(C)** Representative image illustrating classification of 6E10+ plaques as cored or diffuse according to Methoxy labeling (Scale bar, 500µm). Cored plaques were defined as 6E10+, Methoxy+ objects, whereas diffuse plaques were defined as 6E10+, Methoxy-objects. The lower segmentation view shows cored plaques in green and diffuse plaques in red. **(D)** Size distributions of cored and diffuse plaques at 6, 8, 10, and 12 months. The upper plots show object frequency, and the lower plots show percentage of tissue area across plaque-size bins. Corresponding statistical comparisons for all data are presented as heat maps in Supplemental Figures 2 and 3.

The frequency distribution of 6E10+ plaques demonstrated a clear age-dependent shift, with the strongest separation among age groups occurring within the smallest plaque-size bins (Fig. 4B). Small 6E10+ plaques increased progressively from 6 to 12 months, and the 10- and 12-month groups were consistently elevated relative to the 6- and 8-month groups for 6E10+ plaques under ∼2000 µm^2^. In contrast, Methoxy+ plaque and dystrophic neurites showed less pronounced separation across ages. At 6 and 8 months, the frequencies and percent area of total 6E10+ and Methoxy+ plaques were roughly similar, consistent with a balance of compact plaque cores and diffuse plaques at this stage of disease progression. Between 10 and 12 months, however, the abundance of small 6E10+ plaques continued to increase, whereas the corresponding Methoxy+ plaque and dystrophic neurite distributions were almost completely stagnant. Interestingly, the size distribution of dystrophic neurites more closely resembled that of Methoxy+ plaques than 6E10+ plaques. The 10-month group exhibited a higher dystrophic neurite area than the 12-month group in some of the smallest bins and in the bin containing objects larger than 5,000 µm², but these differences did not reach statistical significance after multiple comparisons corrections. Thus, the largest overall shift in plaque frequency and area occurred between the younger, 6- to 8-month animals and the older, 10- to 12-month animals, and for 6E10+ plaques over Methoxy+ plaques or dystrophic neurites.

### Cored plaques dominate at 6 months in the CRND8 model, while small diffuse plaques progress more with age

To better understand how plaque morphology changes with age in this model, we further classified 6E10+ plaques according to Methoxy labeling, with 6E10+Methoxy+ plaques designated as cored and 6E10+Methoxy-plaques designated as diffuse (Fig. 4C). At 6 and 8 months of age, most plaques across the size distribution were cored (Fig. 4D). The frequency and area of cored plaques changed comparatively modestly with age. In contrast, diffuse plaque frequency and area increased markedly with age, particularly within the smallest size bins. Diffuse plaques or plaque clusters larger than 3500 µm² were not detected before 10 months, whereas large cored plaques were present at all ages, albeit at lower frequencies in younger animals.

Taken together, these findings illustrate how plaque pathology accumulates across the examined CRND8 time course, with the most pronounced progression occurring among small diffuse plaques. The continued increase in total plaque burden at later ages appears to be driven primarily by the emergence of new, small diffuse plaques rather than by a proportional increase in Methoxy+ plaque cores. Dystrophic neurite burden closely paralleled the distribution of Methoxy+ plaques, suggesting an association between neuritic damage and plaque core maturation. Collectively, these results indicate that plaque development in CRND8 mice is characterized by a predominance of damaging cored plaques over diffuse plaques in early-mid disease, followed by a phase of rapid accumulation of diffuse plaques and slower development of cores in later disease progression. This suggests that much of the later plaque accumulation in this model consists of non-core-containing depots that may exert distinct pathophysiological effects, underscoring the need to distinguish plaque subtypes when evaluating their relative contributions to synapse loss and functional deficits.

### Plaque morphology and size influence local synapse density profiles

We then combined our plaque population data in Figure 4 with the synaptic imaging around specific populations of plaques from this same experiment. As described for the Rescan filtering, the dataset consisted of well-defined isolated plaques, and plaques with poor definition or clustered plaques were frequently excluded from analysis. These excluded plaques may vary with age group since large plaques and clusters are infrequent and become more common at later ages as shown in Figure 4. However, this strategy was designed to compare synapse density in the most common plaque types across all ages, and our population-level analysis shows that large, diffuse plaques are almost non-existent in our earlier age groups (Fig. 4C). Thus, this approach produces a standardized foundation on which to base subsequent analysis of plaque-associated synapse density across all ages examined.

To determine how plaque characteristics influence local synapse density, HOMER1 density (puncta/µm^2^) was quantified within 6E10+ plaques and associated masks (Fig. 5A). We quantified HOMER1 synapse density in the inner 50% of the 6E10+ area of the plaque, the outer 50% of the plaque, 0-5 µm away, 5-10 µm away, and 10-15 µm away from the plaque. Density was calculated as number of HOMER1 puncta within each mask normalized to the area of each plaque mask. This distance-based analysis provides an estimate of synapse loss within the plaque core relative to the adjacent peri-plaque region. With this method, we quantified baseline synapse density and plaque-associated synapse loss comparable to previous reports from other mouse models. ^7,8,35^

**Figure 5.**
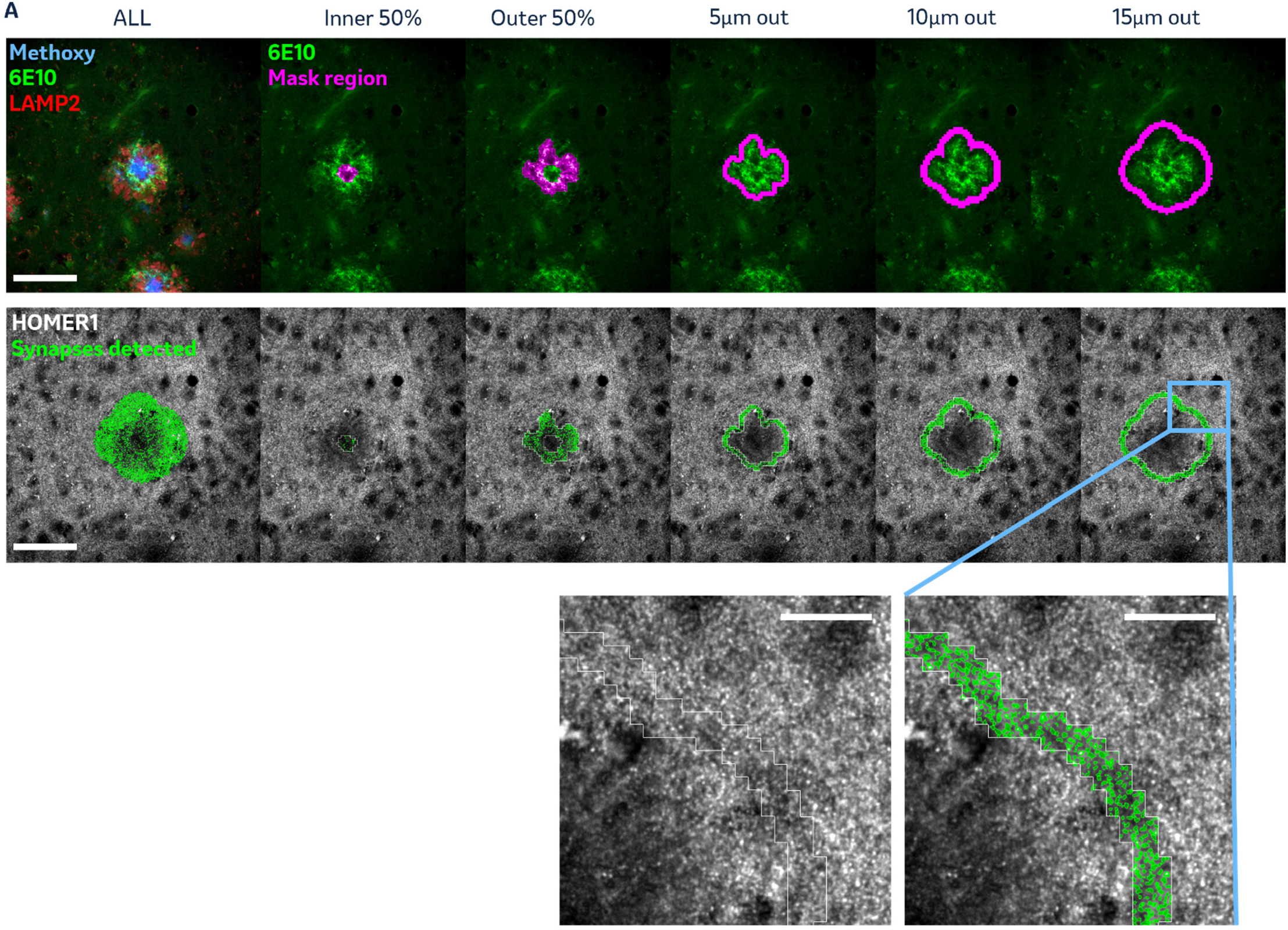
Synaptic puncta detected in plaque-associated masks. **(A)** Representative 63× image and analysis masks used to quantify synapse density as a function of distance from a 6E10+ plaque. The multi-channel raw image in the top left shows a Methoxy+ plaque core surrounded by 6E10+ Aβ and LAMP2+ dystrophic neurites (Scale bars, 50µm). Plaque-associated masks, shown in magenta, divide the 6E10-positive plaque into inner and outer regions and extend outward from the plaque boundary in sequential 5 µm intervals. Lower images show HOMER1 signal and detected synaptic puncta in green for each plaque mask. Enlarged insets illustrate individual puncta detection (Scale bars, 12.5µm).

### Synapse density increases with distance from plaques and is relatively consistent across ages

After post-Rescan filtering of plaques as described in Figure 3, a total of 919 plaques were analyzed across all age groups. When all plaques were analyzed together, synapse density was lowest within the interior of the plaque and increased progressively with distance from the plaque border before reaching a plateau at 5-15 μm from the plaque surface (Fig. 6A). This overall pattern was observed at all ages. When considering age-dependent effects, plaques from older mice surprisingly showed an increased synapse density in the plaque interior compared with younger mice. This effect was also present within only small plaques of 0-1000 μm², as shown in Fig. 6B. Given our findings that aged mice continually accumulate small diffuse plaques at a greater rate than large or cored plaques (Fig. 4), the overall plaque-associated synapse data is likely biased toward higher densities for older animals by small, non-synaptotoxic plaques. Thus, as we have a large dataset, we chose to filter our synapse data by plaque size and morphology across ages to fully understand which plaque features lead to the greatest local synapse loss and therefore contribute most significantly on a tissue-wide scale.

**Figure 6.**
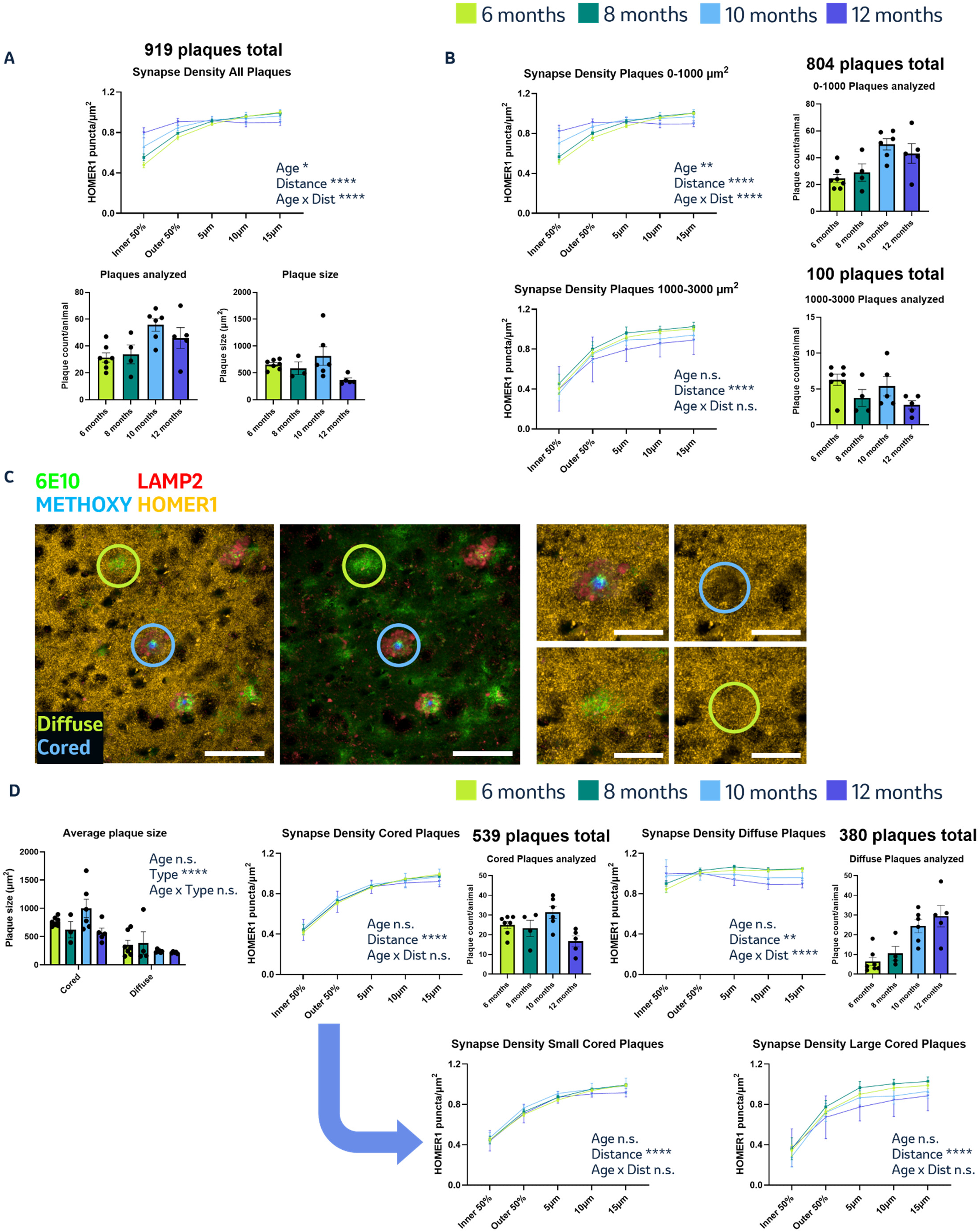
Plaque morphology drives localized synapse loss, while age influences the surrounding tissue. **(A)** HOMER1 synapse density as a function of spatial position relative to all analyzed 6E10+ plaques. A total of 919 plaques were analyzed across the four age groups. The line plot shows synapse density within the inner and outer plaque regions and in masks extending from the plaque boundary. Bar plots summarize the number of plaques analyzed and average plaque size per animal. **(B)** Small plaques were defined as 0-1,000 µm² and included 804 plaques; large plaques were defined as 1,000-3,000 µm² and included 100 plaques. Line plots show spatial synapse-density profiles for small and large plaques, and bar plots summarize the number of plaques analyzed per animal. **(C)** Plaques stratified by core morphology. Representative raw four-channel images show example cored and diffuse plaques within the same imaging field, as indicated by blue and green circles, respectively (Scale bars, 50µm). The inset images on the right indicate visible synapse loss within the region of the cored plaque and relatively normal synapse appearance for the diffuse plaque (Scale bars, 25µm). **(D)** Plaques stratified by core morphology included 539 cored plaques and 380 diffuse plaques. The bar plots summarize the average plaque size per animal within cored and diffuse plaques and the number of plaques analyzed by animal. Line plots show synapse-density profiles across plaque-associated masks. Lower line plots show further stratification of cored plaques into small and large size groups to compare the effects of core status and plaque size on local synapse density. Mixed-effects model main effects and interaction effects are shown on each line graph.

### Larger plaques are associated with greater local synapse depletion

Plaques were filtered by size into small (0-1000 μm²; n = 804 plaques) and large (1000-3000 μm²; n = 100 plaques) groups (Fig. 6B, remaining plaques >3000 um² not included). Small plaques exhibited a moderate reduction in synapse density within the plaque interior, followed by recovery to baseline within 5-15 μm of the plaque boundary. As described above, age-dependent effects for small plaques mirrored those seen in the overall plaque population, in which older animals actually exhibit a higher synapse density within the interior of small plaques. In contrast, large plaques displayed substantially lower synapse density within the plaque interior and a steeper spatial gradient, with synapse density recovering progressively with increasing distance from the plaque surface. No significant age-dependent effect was observed for large plaques.

### Cored plaques induce pronounced local synapse loss, whereas diffuse plaques show minimal local synapse disruption

Plaques were further classified according to morphology using Methoxy labeling to distinguish cored plaques (6E10+ Methoxy+; n=539 plaques) from diffuse plaques (6E10+ Methoxy-; n=380 plaques) (Fig. 6C, D).

Cored plaques exhibited a steep distance-dependent synapse density curve at all ages. Synapse density was lowest within the plaque interior and increased steadily across distance before reaching a plateau approximately 5-15 μm from the plaque surface. The synapse density curves across all age groups were overlapping, and there was no main age effect or interaction effect.

In contrast, diffuse plaques displayed substantially higher synapse density within the plaque interior than cored plaques, and this also led to considerably less spatial variation across distance. Interestingly, similar to that seen for small plaques, age-dependent effects emerged for diffuse plaques, showing higher synapse density in the inner 50% for older animals. However, across the 5-15 μm distances, synapse density decreased with age, especially in the comparisons with 12 month old mice. This points to a more general effect on synapse loss beyond plaques in older animals.

### Plaque core morphology modulates the degree of synapse loss across plaque sizes

As shown in Fig. 4, we observed that the number of diffuse plaques increased progressively with age, whereas the number of cored plaques increased at a slower rate. Although plaque size varied within both populations, comparison of average plaque sizes revealed that the analyzed cored plaques were significantly larger than diffuse plaques at all ages examined (Fig. 6D). Since we have also seen that cored plaques show more localized synapse loss than diffuse plaques, it is possible that our cored vs. diffuse synapse effect is skewed by the average size of the plaques, and that it is the increased area of cored plaques and not the presence of the core that confers toxicity to synapses. As our method produces a large plaque dataset, we were able to further filter cored plaques by the small and large sizes used in Fig. 6B in order to untangle the effects of size and core presence.

Small cored plaques (0-1000 μm²; n = 431 plaques) exhibited a distance-dependent reduction in synapse density that closely mirrored the overall effect for cored plaques (Fig. 6D), once again confirming that small plaques tend to dominate the CRND8 model and the dataset. Large cored plaques (1000-3000 μm²; n = 93 plaques) displayed even lower synapse density within the plaque interior and a steeper recovery gradient than small cored plaques, which mirrors the effect of large plaques. Since we show in Fig. 4D that large plaques are predominantly cored, with several groups showing no diffuse plaques at larger size bins, this suggests that the effects seen in the large size plaques in Fig. 6B are dominated by cored plaques. Thus, cored plaques are the predominant source of local synaptotoxicity, and they can induce this toxicity even at small sizes. Size stratification for diffuse plaques was not performed since no diffuse plaques were detected at many of the large size bins for several groups.

Collectively, these analyses demonstrate that plaque morphology and size are primary determinants of localized synaptic loss, with dense cores and larger plaques driving the most severe synaptic depletion, whereas age exerts comparatively modest influence on plaque-associated synapse loss.

## Discussion

Here we have presented an automated image analysis workflow to enable rapid and scalable characterization of pathological structures across large tissue areas while preserving the spatial and subcellular resolution necessary to interrogate local biological responses. By automating pathology-targeted imaging, this framework substantially increases throughput and minimizes the sampling bias inherent to manual imaging approaches. Furthermore, the ability to stratify individual objects by quantitative morphological features facilitates the investigation of heterogeneous pathological populations, revealing spatial dynamics and potential mechanisms that are masked when all objects are analyzed collectively. By bridging tissue-wide pathology mapping with subcellular measurements, this multiscale framework provides a robust platform to link local biological phenomena to broader disease progression.

The capabilities described here are emerging as part of a broader transformation in biological imaging. New smart acquisition technologies such as ZEISS Guided Acquisition (for Axio and LSM platforms) and Thermo Scientific EurekaScan Finder (for CX7 HCS platforms) integrate automated image analysis with hierarchical imaging workflows to identify structures of interest and prioritize them for targeted characterization. In parallel, artificial intelligence and machine learning approaches are becoming increasingly integrated into imaging platforms, including tools such as the PhenoLOGIC feature of Harmony used in this study, enabling more sophisticated identification and classification of biological structures for downstream analysis. As these tools become implemented across imaging platforms, high-throughput targeted imaging is poised to become widely accessible to investigators studying neurodegenerative disease. This convergence of automation, artificial intelligence, and multiscale imaging has the potential to accelerate discovery and provide new insights into disease mechanisms that are difficult to uncover using traditional imaging approaches.

This automated, multiscale workflow has broad applicability for investigating diverse pathological processes across intact tissues. For example, the field of complement-mediated synaptic pruning in neurodegenerative disease continues to expand and frequently relies on imaging-based approaches to quantify glial engulfment of tagged synapses in intact brain tissue. Beyond synapses, a growing body of evidence implicates additional subcellular and organellar responses, including lipid droplet accumulation, mitochondrial dysfunction, and altered lysosomal processing, as important contributors to disease progression. Furthermore, researchers are increasingly interested in characterizing the local effects of other neurodegenerative protein aggregates, including tau tangles, Lewy bodies, and the complexity of co-pathologies. Quantitatively linking these microscale biological responses to focal lesions across large tissue areas remains technically challenging.

Beyond neurodegenerative disease, this workflow is readily adaptable to other focal pathologies that shape their surrounding microenvironment. These could include infectious lesions such as tuberculosis granulomas or viral infection foci, malignant structures including primary tumors and metastatic lesions, and regenerative or injury-associated sites such as stem cell grafts, stroke lesions, or traumatic tissue damage. Our pathology-centric approach enables quantitative analysis of subcellular biological responses within specific microenvironments while simultaneously capturing the heterogeneity of pathological populations at the tissue-level.

A notable biological finding of this study is that plaque morphology is a stronger determinant of local synaptic vulnerability than plaque size or animal age, with cored plaques exhibiting high local synapse loss and diffuse plaques exhibiting generally low synapse loss. However, across the cortex, the late-stage plaque progression in the CRND8 model was driven largely by the accumulation of small diffuse plaques. It is likely that diffuse and cored plaques represent different stages within a common maturation pathway, but that plaque formation outpaces plaque maturation as disease progresses in this model.

Diffuse plaques have often been interpreted as relatively early Aβ deposits, whereas neuritic or cored plaques are thought to represent more mature lesions characterized by compact fibrillar amyloid, dystrophic neurites, reactive glia, and greater local tissue disruption ^36–38^. Thus, in addition to overall plaque burden, plaque composition and morphology may influence disease progression. This distinction may explain cognitively resilient individuals who tolerate substantial Aβ pathology while exhibiting relatively limited neurodegeneration or cognitive decline. Consistent with this framework, recent evidence suggests that plaque cores develop over time and that increasing plaque age is associated with neuritic dystrophy and synapse loss^39^.

However, the relationship between plaque morphology and toxicity is not explained solely by fibrillar plaque cores. Soluble oligomeric Aβ, which is widely considered the primary synaptotoxic Aβ species, has been detected in association with both diffuse and cored plaques ^40^. Some studies have suggested that oligomeric Aβ creates a toxic halo surrounding plaque cores, which leads to progressive synapse loss as plaques grow into the surrounding tissue ^7,41^. In this context, plaque cores may function less as the direct toxic species and more as structural microenvironments that promote the generation, retention, or concentration of toxic oligomeric Aβ assemblies. The factors that determine why some plaques progress toward a cored state, how plaque maturation is linked to local synaptotoxicity, and the extent to which these processes depend on oligomeric Aβ therefore remain incompletely understood.

Stratifying pathologies to uncover factors that influence disease may hold particular value for human postmortem studies, where disease progression cannot be followed longitudinally. In the present dataset, stratification by plaque size and morphology revealed distinct plaque populations associated with different levels of synaptic injury. Similar approaches could be applied to human tissue to investigate whether morphological or molecularly distinct plaques represent different stages of pathological evolution and whether these states are associated with differential synaptic vulnerability. More broadly, this framework could facilitate efforts to connect specific plaque populations with clinically relevant biomarkers and determine which pathological features are most predictive of neural, and ultimately cognitive outcomes.

In conclusion, this work highlights the power of pathology-centered imaging to move beyond measurements of pathological burden toward understanding pathological heterogeneity and its biological consequences. By linking local microenvironments to tissue-level outcomes, such approaches may help identify the specific pathological features that drive disease progression, therapeutic responses, and cognitive decline.

## Methods

### Animals

Female TgCRND8 mice (Tg(PRNP-APPSweInd)8Dwst, mixed B6;C3H background) from Taconic were aged to 6 (n=7), 8 (n=4), 10 (n=6), and 12 months (n=5). Mice were group housed and given food ad libitum on a 12L:12D light schedule. At necropsy, mice were euthanized with CO^2^ and the body cavity opened for transcardial perfusion of 15-20 mL of PBS. The brain was removed and immediately placed into ice-cold 4% PFA. After 24 hours, brains were washed briefly with PBS and dropped into 30% sucrose in PBS. Brains were stored at 4°C until ready for sectioning. Brains were sent in 30% sucrose to NSA labs for sectioning and staining. All animal procedures were reviewed and approved by the Institutional Animal Care and Use Committee (IACUC) of Merck & Co., Inc., Rahway, NJ, USA (APS 2026-601572-Dec) and were conducted in accordance with applicable institutional guidelines and regulations for the care and use of laboratory animals.

### Neurohistology Embedding, Sectioning & Staining (NSA labs)

Brains received at NeuroScience Associates were examined, then treated overnight with 20% glycerol and 2% dimethylsulfoxide to prevent freeze-artifacts. The specimens were then embedded in a gelatin matrix using MultiBrain® Technology (NeuroScience Associates, Knoxville, TN).

The blocks were rapidly frozen, after curing by immersion in 2-Methylbutane chilled with crushed dry ice and mounted on a freezing stage of an AO 860 sliding microtome. The MultiBrain® blocks were sectioned coronally with 30 micrometer (μ) setting on the microtome. All sections were collected sequentially into series of 24 containers. All containers contained Antigen Preserve solution (50% PBS pH7.0, 50% Ethylene Glycol, 1% Polyvinyl Pyrrolidone); no sections were discarded.

### Immunofluorescence (NSA labs)

For immunofluorescent histochemistry, free floating sections were stained with Methoxy-X04 (Tocris 4920), 6E10 (Novus NBP2-62566), LAMP2 (Abcam ab13524), and HOMER1 (Synaptic Systems 160-019). All incubation solutions from the primary antibody onward use Tris buffered saline (TBS) with Triton X-100 as the vehicle; all rinses were with TBS.

The sections were immunostained with the primary antibodies overnight at room temperature. Vehicle solutions contained Triton X-100 for permeabilization. Following rinses, a species-matched fluoro-tagged antibody from Jackson ImmunoResearch (711-545-152, 712-605-153, 703-165-155) was applied. Following rinses, the sections were mounted on gelatin coated glass slides and air dried. The slides were dehydrated in alcohols, cleared in xylene and coverslipped with Permount.

### Automated high-content imaging and analysis on the Opera Phenix, Harmony

All imaging and object segmentation analysis was performed with Harmony version 5.2.

#### Preciscan overview

The Preciscan function of the Opera Phenix high-content imaging system was used to enable efficient identification and high-magnification imaging of amyloid plaques across MultiBrain block sections. Imaging was focused on the cortex because it accumulates abundant Aβ plaque pathology and a relatively uniform HOMER1-positive synaptic neuropil, enabling consistent automated analysis across large tissue areas. The hippocampus and white matter were excluded because their laminar architecture or low synaptic signal produced large regions with little or no HOMER1 labeling.

A Preciscan Prescan was then performed with a 20× water objective across cortical regions, acquiring multi-channel images suitable for feature-based analysis and plaque selection. Based on these analyses, individual plaques of interest were subsequently imaged with a 63× water objective using the Preciscan Rescan function.

#### Preciscan Prescan set up

MultiBrain slides containing a block section of 22 mouse brains were mounted in a custom 3D-printed holder designed for 2 × 3-inch slides, with the coverslip oriented facing downward. A new slideholder plate type was defined in Harmony including one rectangular “well” for the single 2 × 3-inch slide. Cortical regions of interest were manually selected, with 3 20× fields covering the dorsal cortex. A Preciscan Prescan was configured to acquire a 20× multi-channel z-stack (8 planes, 2 µm step size, NA 1.0, 0.66 µm xy resolution) across cortical regions of interest, including all 4 channels (Methoxy in 405, 6E10 in 488, HOMER1 in 568, LAMP2 in 647) for use in plaque selection and in analysis of plaque and dystrophic neurite populations.

#### Prescan plaque selection

Plaque detection and filtering criteria were designed to select for distinct plaques with sufficient surrounding HOMER1 signal for synapse analysis and to distinguish amyloid plaques from blood vessels. As shown in Figure 2B, we chose to focus imaging on Methoxy+ plaques as Methoxy-diffuse plaques were much more frequent and could still be sampled with Methoxy-focused imaging, whereas Methoxy+ plaques would be undersampled in some groups if all 6E10-positive plaques were included. Plaques sampled included a minimum size of 200 µm^2^ to prevent potential small debris from being imaged and a maximum size of 2500 µm^2^ to avoid poorly-defined plaque clusters. Plaques within 50 µm of regions with low HOMER1 signal (i.e. areas lacking tissue, corpus callosum) were filtered out in order to ensure consistent synaptic signal surrounding selected plaques (Fig. 2B). Blood vessels can exhibit Methoxy positivity due to amyloid beta deposition and therefore were excluded from automatic selection.

We applied two filters taking advantage of blood vessel features: due to their elongated shape, they have a low width:length ratio (cutoff <0.3), unlike plaques which are near circular. Blood vessels also lack dystrophic neurite damage, reflected by low LAMP2 signal, and were also filtered out with a very low LAMP2 intensity threshold. Nearest neighbor distance filtering was applied to exclude plaques with close neighbors, as these would have overlapping domains in downstream synapse analysis. Because synapse loss results in reduced HOMER1 signal at plaque centers, z-planes could not be accurately determined based on plaque center HOMER1 signal, and therefore expanded masks of plaques were created to include surrounding synapse-rich tissue. The resized plaque masks were used for accurate z-plane assignment for each individual plaque based on the plane of maximum HOMER1 signal (Fig. 2B, C). Well layout maps were generated to visualize the x–y positions of selected plaques, and plane maps were used to assign the imaging z-plane for each plaque based on HOMER1 signal (Fig. 2C).

#### Re-scan imaging and analysis

Following X/Y/Z assignment for selected plaques, 63× Rescan imaging was automatically performed. Single-plane images were acquired in all four channels for Methoxy, 6E10, LAMP2, and HOMER1 as described above at 63×, NA 1.15, 0.28 µm xy resolution.

Images were analyzed in Harmony using an automated post-imaging filtering workflow to identify plaques suitable for synapse analysis (Fig. 3A). The 6E10 channel was thresholded to generate the 6E10+ object population. Image regions were then classified using a PhenoLOGIC machine-learning classifier trained to distinguish plaques from non-plaque objects, primarily to exclude blood vessels which can be 6E10+. Objects were retained if they overlapped the PhenoLOGIC plaque-texture class and exceeded a minimum area threshold of 100 µm².

Objects passing the machine-learning and size filters were expanded by 15 µm to create a plaque-associated mask for downstream quality control and synapse quantification. Plaques for which the expanded mask overlapped the image border >2 µm were excluded to avoid incomplete segmentation of plaque-associated masks.

Plaques were also filtered according to nearest-neighbor distance to remove overlapping plaque-associated masks and ensure that synaptic measurements could be assigned to individual plaques. Finally, plaques were filtered on a minimum HOMER1 intensity threshold to ensure reliable detection of synaptic puncta. These quality-control filters generated a final population of isolated plaques with well-defined borders and quantifiable surrounding synaptic signal for downstream analysis.

#### Synapse analysis

For synapse analysis, plaque-associated masks were generated from the final filtered plaque population. Each plaque was subdivided into inner 50% and outer 50% plaque regions, and concentric masks extending 0 to 5 µm, 5 to 10 µm, and 10 to 15 µm beyond the plaque boundary were generated. HOMER1-positive puncta were detected using the Harmony “spots” detection module within the plaque-associated masks.

HOMER1-positive puncta larger than 1 µm^2^ were defined as “debris”, and puncta overlapping these debris objects were excluded from analysis. The remaining puncta were assigned to plaque-associated mask regions and used to calculate synapse density as a function of distance from each plaque.

### Quantification of plaque and dystrophic neurite populations in 20× Prescan images

To quantify cortex-wide pathology populations, 20× Prescan images were analyzed in Harmony using an automated segmentation and classification workflow shown in Supplemental Figure 1. Initial cortical analysis regions were identified from HOMER1 signal. Large regions above a HOMER1 threshold were detected as > 200,000 µm^2^. A PhenoLOGIC machine-learning classifier was trained to distinguish cortex from non-cortical tissue, and large regions with minimal overlap of non-cortex were retained.

Classified cortical regions were consolidated into final cortex masks using distance-based clustering of <100 µm. Cortex region area was recorded for normalization of object population data.

Within cortical masks, objects for population analysis were segmented and quantified. For all objects, a minimum size filter of 50 µm^2^ was applied in order to prevent detection of small image artifacts or debris. 6E10+ plaques were identified by intensity thresholding of the 6E10 channel followed by PhenoLOGIC machine-learning classification as described above for the Prescan plaque selection. Methoxy+ plaque cores were similarly identified by intensity thresholding. As described above, Methoxy+ objects with low LAMP2 signal were filtered out to exclude blood vessel-associated Methoxy signal. Finally, dystrophic neurites were segmented from the LAMP2 channel by intensity thresholding. Area and frequency measurements were recorded for all objects.

To classify plaque morphology, overlap between 6E10+ plaques and Methoxy+ plaque cores was calculated. Plaques exhibiting overlapping (>1%) Methoxy labeling were designated as cored plaques, whereas plaques lacking Methoxy overlap were designated as diffuse plaques.

### Data processing

After exporting plaque and synapse data from Harmony, data was imported and processed in RStudio version 2024.12.0.

#### 20× pathology population data

Harmony-exported object tables were processed in Rstudio to give frequency and area size distributions as seen in Figure 4. Data processing and plaque size-binning analyses were performed using the *tidyverse* package suite for data manipulation and reshaping, including *dplyr* for data wrangling and summarization, *tidyr* for table reshaping, and *stringr* for text processing. Cortical regions were assigned to individual animals and age groups by matching their X/Y positions to a slide map. Tables containing 6E10+ plaques, Methoxy+ plaques, LAMP2+ dystrophic neurites, cored plaques, and diffuse plaques were then linked to the corresponding cortical region and thus assigned back to their animal and age group.

Objects were grouped into 150 µm² area bins from 0 to 5,000 µm², with all objects larger than 5,000 µm² included in a final bin. For each animal and size bin, object number and total object area were calculated. Missing animal-by-bin combinations were assigned a value of zero. Object number per size bin was normalized to the total cortical area to give object frequency/µm², and the summed object area was normalized to the total cortical area to calculate the percent area for each size bin.

Processed data were exported as wide-format tables and combined across all imaged block sections. Values for each animal and object-size bin were averaged across block sections to generate a single mean frequency or percent-area measurement for each size bin per animal. Data averaging was performed using the *tidyverse* package collection for data wrangling, summarization, and file export.

#### 63× synapse data

Data processing was performed using the *tidyverse* package suite for data wrangling, table joining, summarization, reshaping, and file export, and the *purrr* package for iterative merging and transformation of plaque- and animal-level datasets across multiple spot measurement files. At a high level, the processing required a slide map, a plaque table, and spots tables, where each spots table includes spots counted within one plaque mask (e.g. 15µm, 10µm, 5µm, outer 50%, inner 50%). The slide map included a key to indicate the row and column at which each brain was imaged (i.e., Animal 1= 1-1, Animal 2= 1-2, etc.). The plaque table contained one row per plaque and included plaque metadata (image field, object number, slide coordinates) as well as the area measurements for the masks used to quantify synapse spots (15µm area, 10µm area, 5µm area, outer 50% area, inner 50% area). Each plaque was assigned to an animal and treatment by inferring the plaque’s row and column number from its slide coordinates and then was matched to the slide map to get the animal and treatment.

For each spots table, the plaque object number column was identified in order to link each spot back to a plaque. Each spot was matched with a plaque and inherited the corresponding metadata. Duplicate measurements that arise from overlapping image fields were then removed: a single plaque-associated spot can sometimes be detected in more than one imaging field when nearby plaques are imaged, and thus each image field where the spot was measured was compared to the preselected field for that plaque in the plaque table. Only spots from the designated field were retained. Once the correct spots were identified, the appropriate area column was determined from the plaque table (e.g., 5µm spots use the 5µm area, inner 50% spots use the inner 50% area, etc.), and each spot was matched to its intended mask area measurement. This allowed calculation of synapse density as number of spots in each mask normalized to the corresponding plaque mask area for each plaque.

Finally, the cleaned data was summarized at two levels. First, plaque-level spot density was calculated for every plaque mask. Spot densities were then averaged for all plaque masks across each animal and merged into a single table. Data averaging was performed using the *tidyverse* package collection for data wrangling, summarization, and file export.

## Statistical analysis

All statistical analyses were run in Graphpad Prism version 10.2.2. Heatmaps were generated in RStudio version 2024.12.0 using *tidyverse* for data wrangling with *ggplot2* for heatmap plots.

For object population data in Figure 4, since each mouse contributed frequency and percent area measurements across many size bins, data were analyzed using a mixed-effects model. Age group and plaque area bin were included as fixed effects, while subject was included as a random effect to account for correlations among measurements obtained from the same animal. A full factorial model including the age × plaque area bin interaction was fitted. Post hoc pairwise comparisons between age groups were performed within each plaque-area bin using Tukey’s multiple-comparisons test.

For plaque size comparisons between cored and diffuse plaques in Figure 6, a two-way ANOVA with main effects of age and plaque type was used. Post hoc pairwise comparisons between age groups were performed within each plaque type using Tukey’s multiple-comparisons test.

For synapse density data in Figure 6, a mixed-effects model was used to accommodate repeated measurements within subjects. Age group and distance from plaques were included as fixed effects, while subject was included as a random effect to account for correlations among measurements obtained from the same animal. A full factorial model including the age × plaque distance interaction was fitted. Post hoc pairwise comparisons between age groups were performed within each plaque distance mask using Tukey’s multiple-comparisons test.

## Author Contributions

Method design: C.A.P.; Experimentation: C.A.P., M.H.; Analysis and Coding: C.A.P., R.P.; Project supervision: X.W., S. S., R.K., M.M.; Manuscript preparation: All

## Competing Interests

At the time this research was conducted, all authors were employees of Merck Sharp & Dohme LLC, a subsidiary of Merck & Co., Inc., Rahway, NJ, USA, the sponsor of the research.

## Acknowledgements

The authors acknowledge support from the MRL Postdoctoral Research Program. The authors thank the 3D Technologies team at our West Point site for assistance designing the custom slideholder used throughout the imaging experiments. The authors thank Will Grever at Revvity for expert technical guidance and support in developing and optimizing the Opera Phenix Preciscan workflow used in this study. The authors thank Neuroscience Associates for the high quality of their services.

