## Supplemental figures for "Automated multiscale high-content imaging connects plaque population heterogeneity to localized synapse loss"

#### Supplemental Figure 1. Prescan image analysis for Methoxy, 6E10 and DNs population data

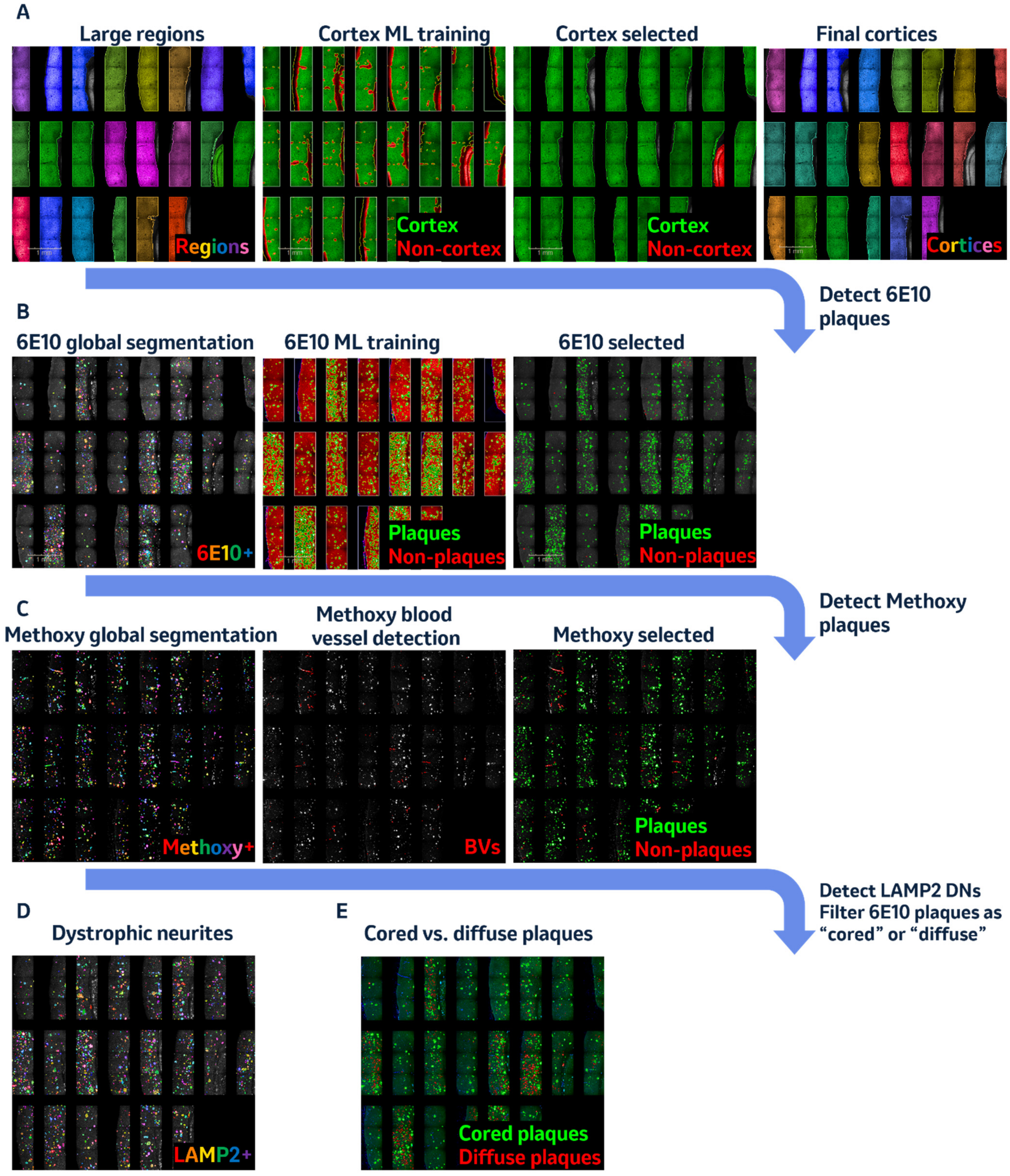

**Supplemental Figure 1. Prescan image analysis for population data**

**(A)** Identification of cortical tissue regions. Large tissue regions were initially detected, a PhenoLOGIC machine-learning classifier was trained to distinguish cortex from non-cortical tissue, and selected cortical regions were combined to generate the final cortical analysis masks. Colors distinguish separate brains in the representative whole-slide views. **(B)** Detection of 6E10+ plaques. Global 6E10 thresholding was followed by PhenoLOGIC machine-learning training and selection of objects classified as plaques, with the same selection process as described for Figure 3. **(C)** Detection of Methoxy+ plaque cores. Global Methoxy thresholding was followed by identification and exclusion of Methoxy-positive blood-vessel signal (Methoxy$^{hi}$, LAMP2$^{lo}$), producing the final set of selected Methoxy+ plaque objects. **(D)** LAMP2+ objects were detected as dystrophic neurites (DNs). **(E)** 6E10+ plaques were classified as cored or diffuse according to the presence or absence of overlapping Methoxy labeling, respectively. The resulting object populations were used for tissue-wide quantification of object frequency and percentage of cortical area.

#### Supplemental Figure 2. Statistical comparisons of 6E10 plaques, Methoxy plaques, and LAMP2 dystrophic neurites across age and size bins

**A** 6E10 frequency

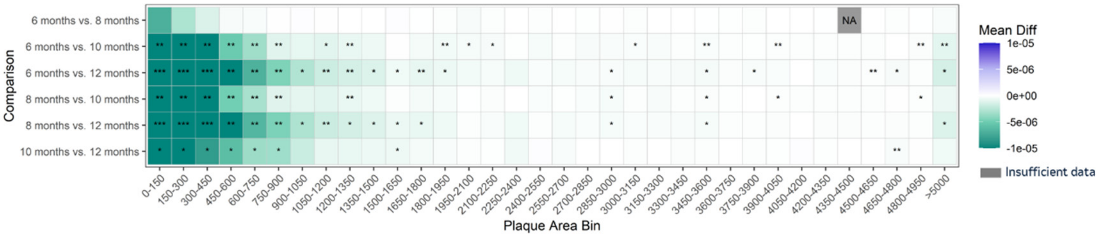


6E10 area

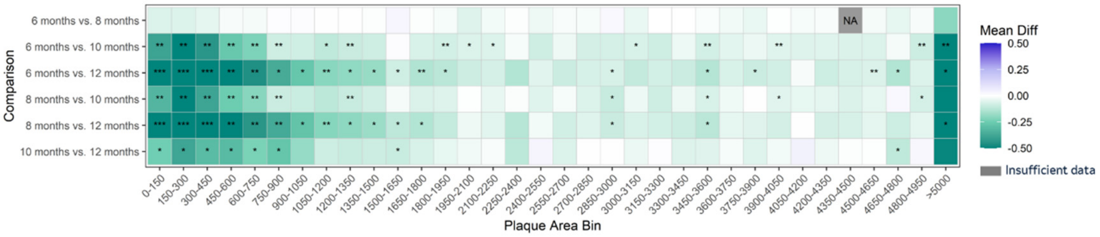


**B** Methoxy frequency

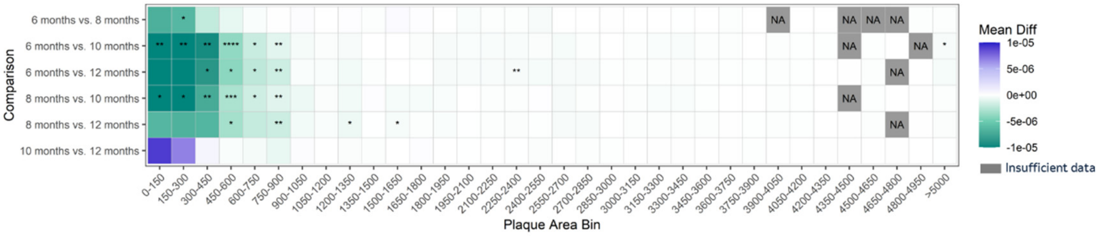


Methoxy area

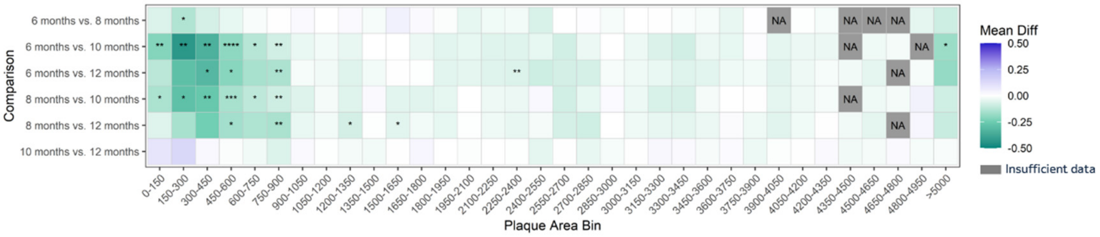


**C** DNs frequency

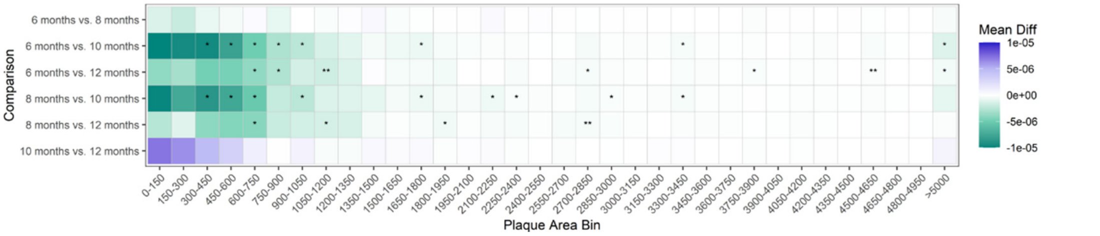


DNs area

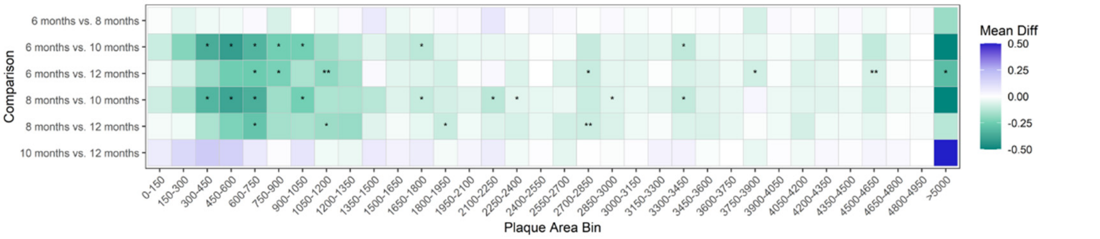

**Supplemental Figure 2. Statistical comparisons of 6E10 plaques, Methoxy plaques, and dystrophic neurites across age and size bins**

Heatmaps summarize object population mixed-effects model statistics with Tukey's multiple comparisons at each size bin across age groups for **(A)** 6E10 plaque frequency and percent area, **(B)** Methoxy plaque frequency and percent area, and **(C)** DNs frequency, and DNs percent area. Color intensity represents the absolute mean difference according to the scales shown, with teal representing an increase in values with age and blue representing a decrease. * adj $p < 0.05$, ** adj $p < 0.01$, *** adj $p < 0.001$, **** adj $p < 0.0001$. NA indicates invalid comparisons for which there were insufficient data.

### Supplemental Figure 3. Statistical comparisons of cored and diffuse plaque populations across age and size bins

## A

##### Cored plaque frequency

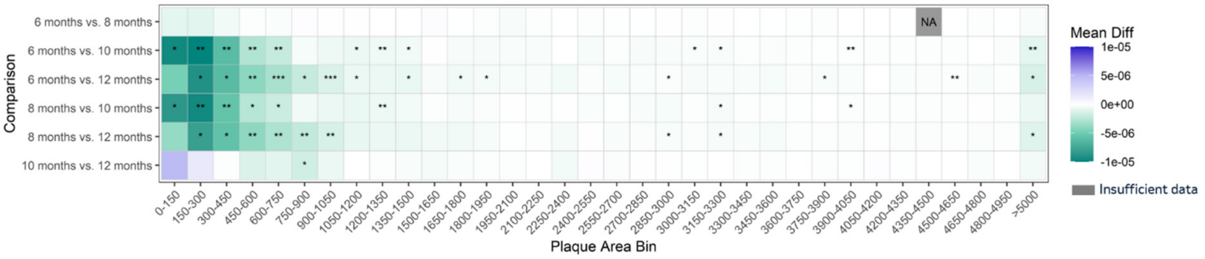


##### Cored plaque area

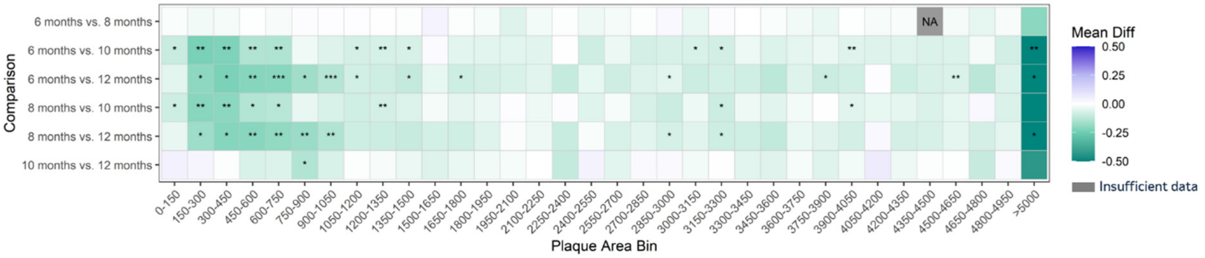


## B

##### Diffuse plaque frequency

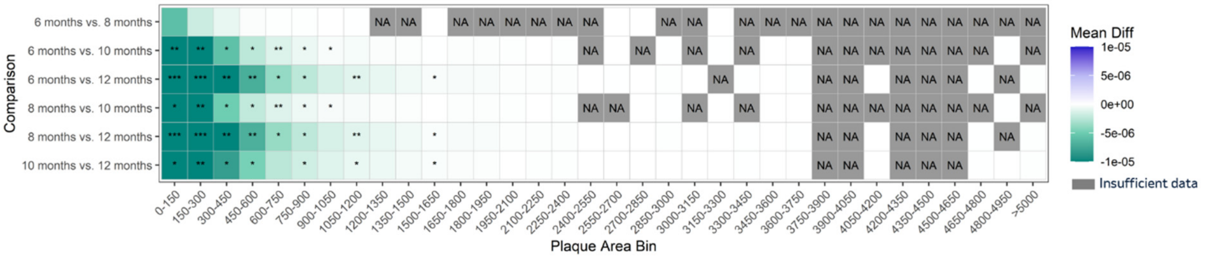


##### Diffuse plaque area

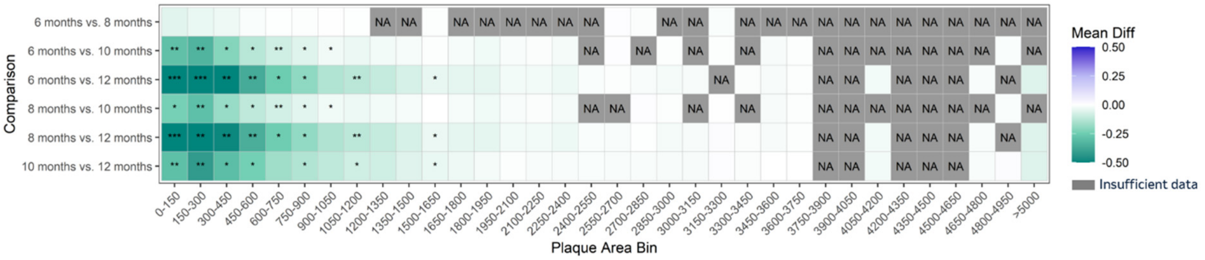

**Supplemental Figure 3. Statistical comparisons of cored and diffuse plaque populations across age and size bins**

Heatmaps summarize object population mixed-effects model statistics with Tukey's multiple comparisons at each size bin across age groups for **(A)** cored plaque frequency and percent area and **(B)** diffuse plaque frequency and percent area. Cored plaques were defined as 6E10+, Methoxy+ objects, and diffuse plaques were defined as 6E10+, Methoxy- objects. Color intensity represents the absolute mean difference according to the scales shown, with teal representing an increase in values with age and blue representing a decrease. * adj $p < 0.05$, **adj $p < 0.01$, *** adj $p < 0.001$, **** adj $p < 0.0001$. NA indicates invalid comparisons for which there were insufficient data.

#### Supplemental Figure 4. Statistical comparisons of synapse density across age, plaque type, and distance from plaque

**A**

##### Synapse Density All Plaques

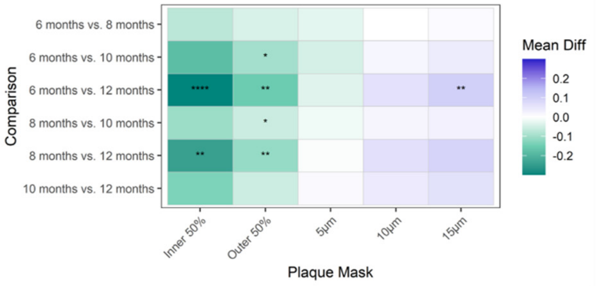


**B**

##### Synapse Density 0-1000μm²

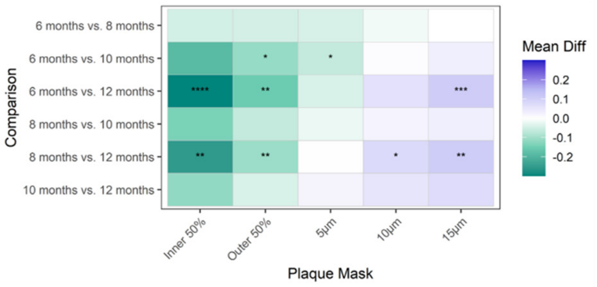


##### Synapse Density 1000-3000μm²

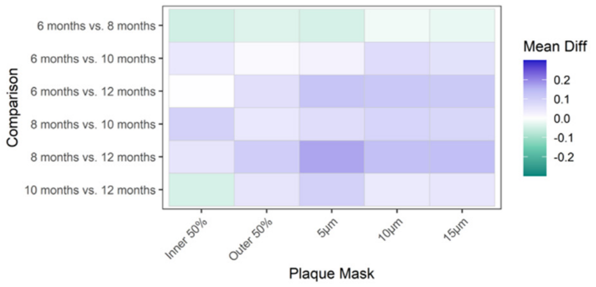


**C**

##### Synapse Density Cored Plaques

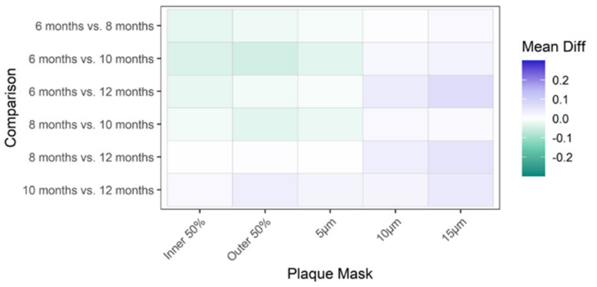


##### Synapse Density Diffuse Plaques

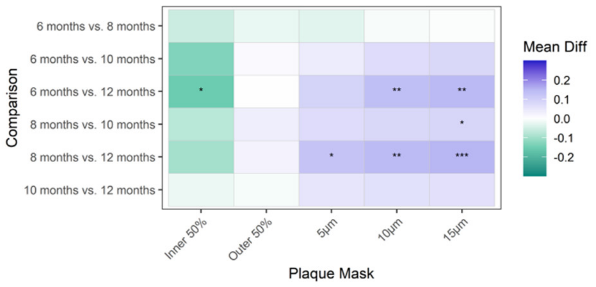


**D**

##### Synapse Density Small Cored Plaques

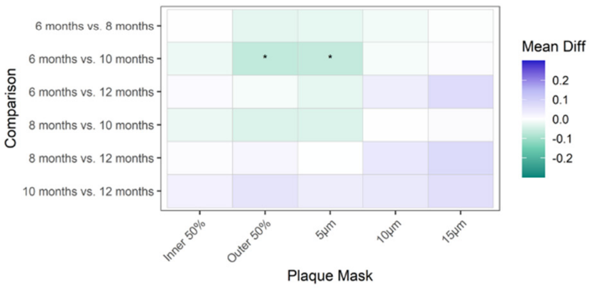


##### Synapse Density Large Cored Plaques

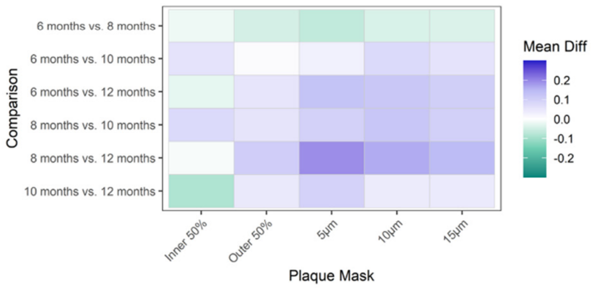

**Supplemental Figure 4. Statistical comparisons of synapse density across age, plaque type, and distance from plaque**

Heatmaps summarize synapse density mixed-effects model statistics with Tukey's multiple comparisons at each distance from plaque across age groups for **(A)** all plaques, **(B)** small and large plaques, **(C)** cored and diffuse plaques, and **(D)** small and large cored plaques. Color intensity represents the absolute mean difference according to the scales shown, with teal representing an increase in values with age and blue representing a decrease. * adj $p < 0.05$, **adj $p < 0.01$, *** adj $p < 0.001$, **** adj $p < 0.0001$.
